# The impact of Quaternary Amazonian river dynamics on patterns and process of diversification in uakari monkeys (genus *Cacajao*)

**DOI:** 10.1101/2023.06.23.546215

**Authors:** Felipe Ennes Silva, Leilton Willians Luna, Romina Batista, Fabio Röhe, Chrysoula Gubili, Izeni P. Farias, Tomas Hrbek, João Valsecchi do Amaral, Camila C. Ribas, Allan D. McDevitt, Simon Dellicour, Jean-François Flot, Jean P. Boubli

## Abstract

**Aim:** Western Amazonia is a region that underwent several landscape changes during the Quaternary. While Riverine Barrier Hypothesis is traditionally used to explain the influence of rivers on speciation, processes such as river rearrangements have been overlooked to explain the geographic distribution and evolutionary history of the Amazonia biota. Here we test how river rearrangements in western Amazonia influenced the evolutionary history of uakari monkeys, a primate group most associated with seasonally flooded forests in western Amazonia.

**Location:** Western Amazonia

**Taxon:** The uakari monkey (genus *Cacajao*)

**Methods:** We performed a continuous phylogeographic analysis using 77 cytochrome *b* sequences and used digital elevation models to identify the role of landscape and riverscape characteristics in the geographic distribution of *Cacajao*. Finally, we used genome-wide SNPs variation (ddRADseq) to investigate population structure, gene flow and demographic history in three *Cacajao* species that were impacted by river rearrangements.

**Results:** Our continuous phylogeographical reconstruction points that the ancestral *Cacajao* lineage occupied the flooded forests of the Solimões River at ∼1.7 Mya, and descendant lineages dispersed throughout western Amazonia more recently. We identified gene flow among both black and bald-headed uakari populations, even across rivers considered barriers (e.g., the Negro River). Landscape analysis showed that river rearrangements influenced the geographic distribution and population structure in *Cacajao*. The demographic analysis indicates that *C. calvus, C. amuna*, and *C. rubicundus* went through a population decline in the last 70 Kya and have a low effective population size.

**Main conclusion:** Our results support that the river rearrangements have shaped the geographic distribution and divergence of recently diverged *Cacajao* lineages. Landscape and riverscape changes, along with retractions of the flooded forests, isolated some *Cacajao* populations in floodplain areas. Our study also suggests that these events led to the recent population decline in species with a restricted geographic distribution.

## INTRODUCTION

The Amazon Rainforest hosts an exceptionally high biodiversity that evolved through an intricate interplay between abiotic and biotic factors (Antonelli et al., 2018, Cracraft et al., 2020). Eastern and western Amazonia have different geomorphological and climatic dynamics that have shaped biological evolution in various ecological systems (Oberdorff et al., 2019; Silva et al., 2019, Miranda et al., 2022). In eastern Amazonia, the Guiana and Brazilian shields are older and geologically more stable, while In the central and western Amazonia, the intense sedimentological and tectonic activity promoted changes in the river network during the Quaternary, including avulsions, incisions and river capture events (Rossetti, 2014, Ruokolainen et al., 2019, Pupim et al., 2019, Rossetti et al., 2021). This intense river dynamics consequently affected extensive areas of seasonally flooded forests and had an impact on the diversification of species inhabiting these environments (Thom et al., 2020; Luna et al., 2022, 2023). However, the impact of these landscape dynamics on the geographic distribution and demographic history of floodplain-adapted species in western Amazonia remains fairly unexplored (but see Barbosa et al., 2022).

Evolutionary studies of primates have played a major role in understanding Amazonian biogeography. Since Wallace’s seminal work (Wallace, 1852) that inspired the Riverine Barrier Hypothesis (RBH, Gascon et al., 2000), new information on primate diversity and distribution has allowed us to refine our understanding of the role of Amazonian rivers and its associated habitats as drives of speciation in this group (Ayres & Clutton-Brock, 1992, Boubli et al., 2015, Fordham et al., 2020, Janiak et al., 2022, Mouthé et al., 2022). General findings indicate that Amazonian rivers – traditionally considered geographic barriers to gene flow – can be effective in keeping apart some populations and species of small-body primates such as titi monkeys, tamarins and marmosets, but are relatively more permeable to many medium and large-body primates (Janiak et al., 2022, Mourthé et al., 2022). The influence of river rearrangements in the diversification of Amazonian primates was never studied, although it can play a major impact in the evolutionary history of those species occurring in the floodplain areas of western Amazonia.

*Cacajao* is a genus with distribution limited to western Amazonia, while its sister group, the genus *Chiropotes*, occupy the upland forests of the Guiana and Brazilian shields in eastern Amazonia (Silva Jr et al., 2013). The two groups diverged by approximately 5.1 Mya (Silva et al., 2022). Species-level diversification in *Cacajao* has occurred since 0.5 Mya and may have been influenced by rivers and flooded forest rearrangements in western Amazonia (Silva et al., 2022). The genus comprises eight species currently recognised, including three species of black uakaris (*C. ayresi, C. hosomi*, and *C. melanocephalus*, Boubli et al., 2008) and five species of bald-headed uakaris (*C. amuna, C. calvus, C. rubicundus, C. ucayalii*, and *C. novaesi* Silva et al., 2022). While the eight *Cacajao* species were well-delimited lineages in a genome-wide phylogeny, bald-headed uakaris are separated into only two main lineages if the mitochondrial cytochrome *b* gene is used for phylogenetic inference (Silva et al., 2022).

Seasonality is a critical variable in the ecology of uakaris, affecting their habitat use in response to food availability (Barnett et al., 2013). Therefore, uakaris may either seasonally migrate between flooded forests (*várzeas* and *igapós*), non-flooded forests (*terra firme*), and palm swamps, or live permanently in these habitats (Barnett et al., 2013). An important aspect to investigate is if they live permanently in a certain habitat because they are finding all requirements, or if they are there because of the lack of connectivity to other habitat types (or if both situations operate simultaneously). For example, *C. calvus* has a population living entirely in the flooded forests of the Mamirauá Sustainable Development Reserve (Mamirauá SDR), which is limited by large rivers such as the Solimões and Japurá (Silva et al., 2021). Likewise, some populations of the Peruvian red uakaris (*C. ucayalii*) have been reported living in altitude areas (>600 m a.s.l.), completely isolated from floodplain forests (Heymann & Aquino, 2010; Vermeer et al., 2013, Silva et al., 2021). These populations became isolated from others and disjunct distributions have been reported (Veermer et al., 2013, see also MacHugh et al., 2019, supplementary material). However, historical factors such as landscape changes have constrained the distribution of some populations in Brazil, limiting them to floodplain areas.

The region where bald-headed uakaris occur has undergone intense landscape and riverscape changes throughout the Quaternary (Ruokolainen et al., 2020, Rossetti et al., 2021). Furthermore, in central and western Amazonia, sedimentary deposits followed by regional channel incisions caused flooded forests to retract and upland forests to expand in the late Pleistocene (particularly within the last 200 thousand years ago; kya) (Pupim et al., 2019). This dynamic likely promoted periodic range contractions and isolation of bald-headed uakari populations, possibly explaining the current disjunct distribution of *C. calvus* and *C. rubicundus*. These changes in the landscape may have facilitated successive gene flow episodes between populations, as evidenced by different tree topologies observed in nuclear and mitochondrial datasets (Silva et al., 2022). Therefore, population differentiation will depend on the species’ dispersal capacity, habitat use, and adaptive potential, constrained by landscape features and environmental conditions of these newly formed areas.

In this study, we addressed the following question: how have landscape dynamics in western Amazonia influenced the geographic distribution, demographic history, and diversification of *Cacajao*? We approached this issue by (1) performing a Bayesian phylogeographic analysis to estimate the centre of origin of the ancestral *Cacajao* lineages and their dispersion in western Amazonia; (2) investigating the association between Amazonian river locations and divergence time alongside landscape and riverscape; (3) assessing genome-wide levels of genetic diversity and gene flow across all *Cacajao* species using reduced-representation sequencing; and (4) exploring the demographic history of three closely related uakari species (*C. amuna, C. calvus*, and *C. rubicundus*) in the Solimões, Jutaí, and Juruá river basins, a region where evidence of river position shifts and their impact on flooded and non-flooded forests have been presented (Pupim et al., 2019, Ruokolainen et al., 2020, Rossetti et al., 2021). If the seasonally flooded forests are an essential habitat for species, we would expect that the retraction of the flooded forests potentially led to population declines. Investigating the role of these dynamics in population contact and isolation, expansions and contractions, will enhance our understanding of the evolutionary history of Amazonian species and the exceptional diversity of this biome.

## METHODS

### Divergence time and phylogeographic analyses

We used 77 cytochrome *b* sequences available in GenBank (Table S1) to estimate the divergence time of the *Cacajao* lineages by performing a time-scaled phylogenetic inference using the software package BEAST 1.10.4 (Suchard et al., 2018). We included two *Chiropotes albinasus*, a *Pithecia irrorata* and two titi monkeys – *Plecturocebus cupreus* and *Cheracebus purinus* – as outgroups and calibrated the tree using the age estimated for the fossil genus *Cebupithecia*, the oldest known pitheciin fossil (Beck et al., 2023), timing the divergence between Callicebinae/Pitheciinae. We aligned the sequences using the Mafft online server (Katoh et al., 2019) under the iterative refinement option (FFT-NS-1). This option improves the alignment accuracy for a small number of sequences (Katoh et al., 2019). We modelled the nucleotide substitution process according to an HKY+Γ parameterisation – identified by the ModelFinder algorithm (Kalyaanamoorthy et al., 2017) – as the best substitution model, and the branch-specific evolutionary rates according to a relaxed molecular clock with an underlying log-normal distribution. We used the Bayesian skyline coalescence model (Gernhard et al., 2008) for the tree topology, following Silva et al., (2022). We ran the Markov chain Monte Carlo (MCMC) for 200,000,000 generations, sampling every 50,000 steps, and visually assessed the convergence, mixing, and 10% burn-in using the program Tracer 1.7 (Rambaut et al., 2018).

To perform a continuous phylogeographic analysis, we used the relaxed random walk diffusion model implemented in the Bayesian framework of the software package BEAST 1.10.4 (Lemey et al., 2010). This model has been applied to investigate vertebrates’ phylogeography and evolutionary history (Camargo et al., 2013; Lynch Alfaro et al., 2015; Nascimento et al., 2013; Werneck et al., 2015). For our continuous phylogeographic analysis, we used a log-normal distribution to model the among-branch heterogeneity in diffusion velocity (Lemey et al., 2010, Pybus et al., 2012). Considering that some specimens were assigned to the exact same sampling coordinates, we used the option “add random jitter to tips” and set the values to 0.50 to get slightly different coordinates assigned to the tips of the tree. We used the R package “seraphim” (Dellicour et al., 2016a, 2016b) to extract the temporal information embedded within posterior trees and to visualise the continuous phylogeographic reconstruction.

### Landscape changes in western Amazonia, and the geographic distribution of uakari monkeys

Digital elevation models (DEM) from the Shuttle Radar Topography Mission (SRTM) have been used to identify and characterize the paleochannels in western Amazonia (Ruokolainen et al., 2020; Rossetti et al., 2021). We plotted the locality points of uakaris in the DEM-SRTM image and looked for topographical patterns in the river floodplains – river captures, avulsions, and tributary rearrangements – to explore the effects of landscape and riverscape changes in the current pattern of uakaris’ geographic distribution. This analysis allows us to identify historical landscape and riverscape changes that may have promoted the contact and isolation of uakari populations. We used the software QGIS v. 3.24.2 (QGIS Development Team, 2021) to analyse the images and produce the output map.

### Population genetic variation in *Cacajao*

We used 47 *Cacajao* sequences produced from ddRADseq (Peterson et al., 2012) available in the NCBI database (https://www.ncbi.nlm.nih.gov/; Table S1). We assessed the quality of the raw ddRAD data using FastQC 0.11.8 (Andrews, 2018) and trimmed the reads to 250 bp using the program STACKS v.2.4 (Catchen et al., 2011, 2013; Rochette et al., 2019). We kept only those reads with PHRED scores > 30 for the subsequent analyses. We used *Pithecia pithecia* (GenBank: PVIP00000000.1) as the reference genome in the ipyrad software pipeline (Eaton and Overcast, 2020) to select our ddRAD loci and SNPs. We set the ipyrad “*minimum samples per locus*” parameter to 12, keeping only loci present in at least ∼25% of our sequences. As a result, we retrieved 9,303 loci in a matrix with 2,287,515 positions (33.62% missing sites) and 58,791 SNPs in a matrix with 30.80% of missing sites.

We performed a principal component analysis (PCA) to summarise the genetic variation among *Cacajao* species and populations. We first ran a PCA analysis using all *Cacajao* samples and, after that, we ran a PCA separately for black (*C. melanocephalus, C. ayresi, C. hosomi*) and bald-headed (*C. calvus, C. amuna, C. rubicundus, C. ucayalii, C. novaesi*) uakaris. Populations were identified based on their geographic distribution (Figure S1; Table S2).

We tested shared genetic variation in *Cacajao* species using the ancestry graph model approach implemented in the software package TreeMix (Pickrell & Pritchard, 2012). We used a matrix with one SNP per locus to reduce the effect of linkage and ran four iterations for black uakaris and for bald-headed uakaris. After filtering, we got 6,220 unlinked SNPs across 17 samples of black uakaris and 3,673 unlinked SNPs across 26 samples of bald-headed uakaris.

### Demographic analyses

We tested alternative demographic models using the program FASTSIMCOAL 2.5.2 (FSC - Excoffier et al., 2013) to investigate divergence processes among *C. rubicundus, C. calvus*, and *C. amuna*. These species occur in a region where river positions have shifted in the last 200 kya (see (Pupim et al., 2019, Ruokolainen et al., 2020, Rossetti et al., 2021) and impacted their predominantly flooded forest habitats. Consequently, processes such as gene flow and population size may have been influenced by landscape changes and habitat availability.

With this simulation-based approach, we inferred parameters of divergence, migration, and changes in effective population size from the likelihood of the joint Site Frequency Spectrum (jSFS) obtained from the SNPs datasets of the three taxa calculated in dadi 1.7 (Gutenkunst et al., 2009). We used information on the phylogenetic relationship, gene flow, and population structure information to design and compare four demographic models. These models were then linked to different proposed scenarios of Amazon landscape evolution during the Pleistocene (Pupim et al., 2019; Ruokolainen et al., 2020; Rossetti et al., 2021). The proposed demographic models incorporated hypothetical scenarios in which the isolation of taxa has resulted from the retraction of floodplain forests and the expansion of terra-firme forests, potentially diminishing gene flow and inducing population declines. The best-fit model was selected in a hierarchical approach, first accounting for the reduced presence of gene flow during the divergence of the three taxa, with Model (1) assuming isolation with gene flow among the three taxa, and Model (2) considering isolation with gene flow only between *C. calvus* and *C. amuna*, but the absence of gene flow between *C. rubicundus* and *C. calvus*. These models were designed based on the combined evidence of gene flow between well-delimited populations. We then selected the best model and tested different post-divergence population syndromes, described here as Model (3), a scenario of exponential population expansion due to increasing areas with suitable habitats for these taxa (Barbosa et al., 2022; Sawakuchi et al., 2022). Alternatively, we simulated exponential population decline in Model (4), considering a scenario of reduced habitats, mainly seasonally flooded forests, during the recent past (Pupim et al., 2019).

To select the best-fitting model, we compared the runs with the highest composite likelihood for each model by evaluating the relative weight of the Akaike Information Criterion (AIC - Akaike, 1974). For parameter estimates and model comparison, we performed 100 runs for each model with 50 conditional maximization algorithm cycles and 100,000 simulations for probability maximization. The demographic parameters of the best-fit model, as well as the confidence intervals, were calculated from 100 parametric bootstrap replications, using jSFS obtained from the simulation run with the best composite likelihood value. For time estimates, we applied a mutation rate of 8.48 x 10^−9^ substitutions per site per generation (Kuderna et al., 2023).

## RESULTS

### Divergence time and phylogeographic analyses

The Pitheciinae clade diverged from the clade of titi monkeys – *Plecturocebus, Cheracebus*, and *Callicebus* – at 14.7 Mya (95% highest posterior density [HPD]: 13.87 – 16.08 Mya). The divergence time between *Pithecia* and *Chiropotes*/*Cacajao* occurred at 7.12 Mya (95% HPD: 4.18 – 12.54 Mya), with the split between *Chiropotes* and *Cacajao* estimated at 4.83 Mya (95% HPD: 3.88 – 5.8 Mya).

The continuous phylogeographical reconstruction points to the flooded forests of the Solimões River, central Amazonia, as a geographic origin of the *Cacajao* at ∼1.7 Mya (Figure 1). At ∼0.7 Mya, the ancestral lineage of black uakaris occupied the Negro-Japurá interfluve, to the North; while the ancestral lineage of bald-headed uakaris occupied the southern bank of the Solimões River basin, and the Jutaí River basin, to the South (Figure 1). The diversification in *Cacajao* started between 0.7 and 0.4 My. From 0.5 Mya, the black uakaris’ lineages dispersed toward the North of Brazil, nearby the border with Colombia and Venezuela, while the bald-headed uakaris’ lineages dispersed toward the upper Juruá River and throughout the Javari and Ucayali River basin. From <0.1 Mya, *Cacajao* lineages were spread through their current known geographic range (Figure 1).

**FIGURE 1.**
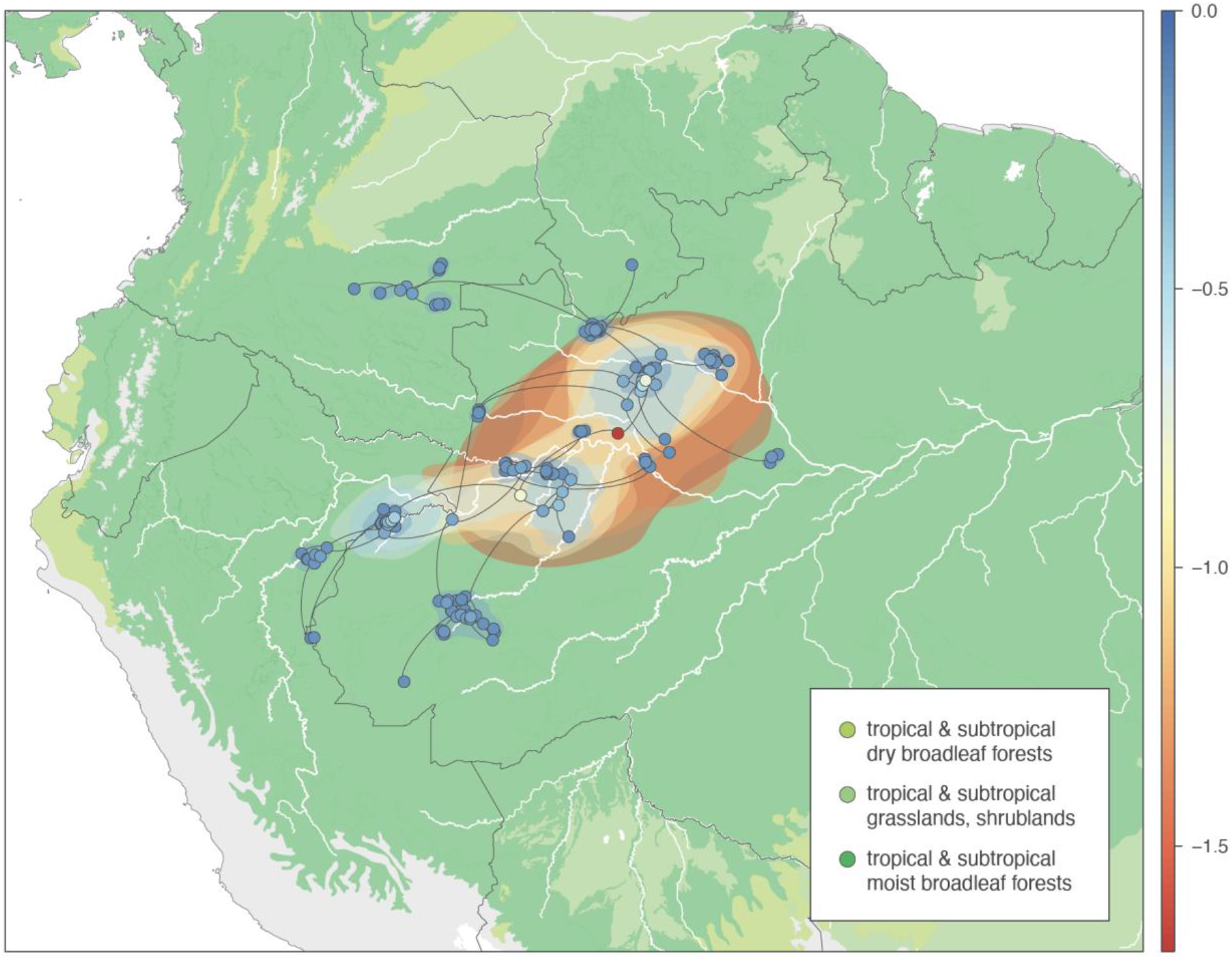
Continuous phylogeographic reconstruction of the dispersal history of *Cacajao* lineages. We mapped the maximum clade credibility (MCC) tree whose nodes are coloured from red (the time of the most recent common ancestor, TMRCA) to blue (the most recent sampling time). This MCC tree is superimposed on 80% highest posterior density (HPD) polygons reflecting phylogeographic uncertainty and coloured according to the same time scale (in Mya).

### Landscape changes in western Amazonia, and the geographic distribution of uakari monkeys

We identified topographic features of fluvial origin related to river rearrangements that coincide with the geographic separation of *C. calvus* and *C. rubicundus* populations (Figure 2). The westward displacement of the Juruá River by avulsion during the Pleistocene-Holocene left behind a huge paleo valley. In the DEM-SRTM images, is possible to identify a paleochannel that connected the middle Juruá River to the lower Jutaí River, which is currently a floodplain limited by the Jutaí and Riozinho rivers and occupied by a population of *C. calvus* (Figure 2, see also population calJT, Fig. S1 and Table S2). This population is separated by about 100 km from the population of *C. calvus* that occurs in the flooded forests of the Mamirauá Sustainable Development Reserve.

**FIGURE 2.**
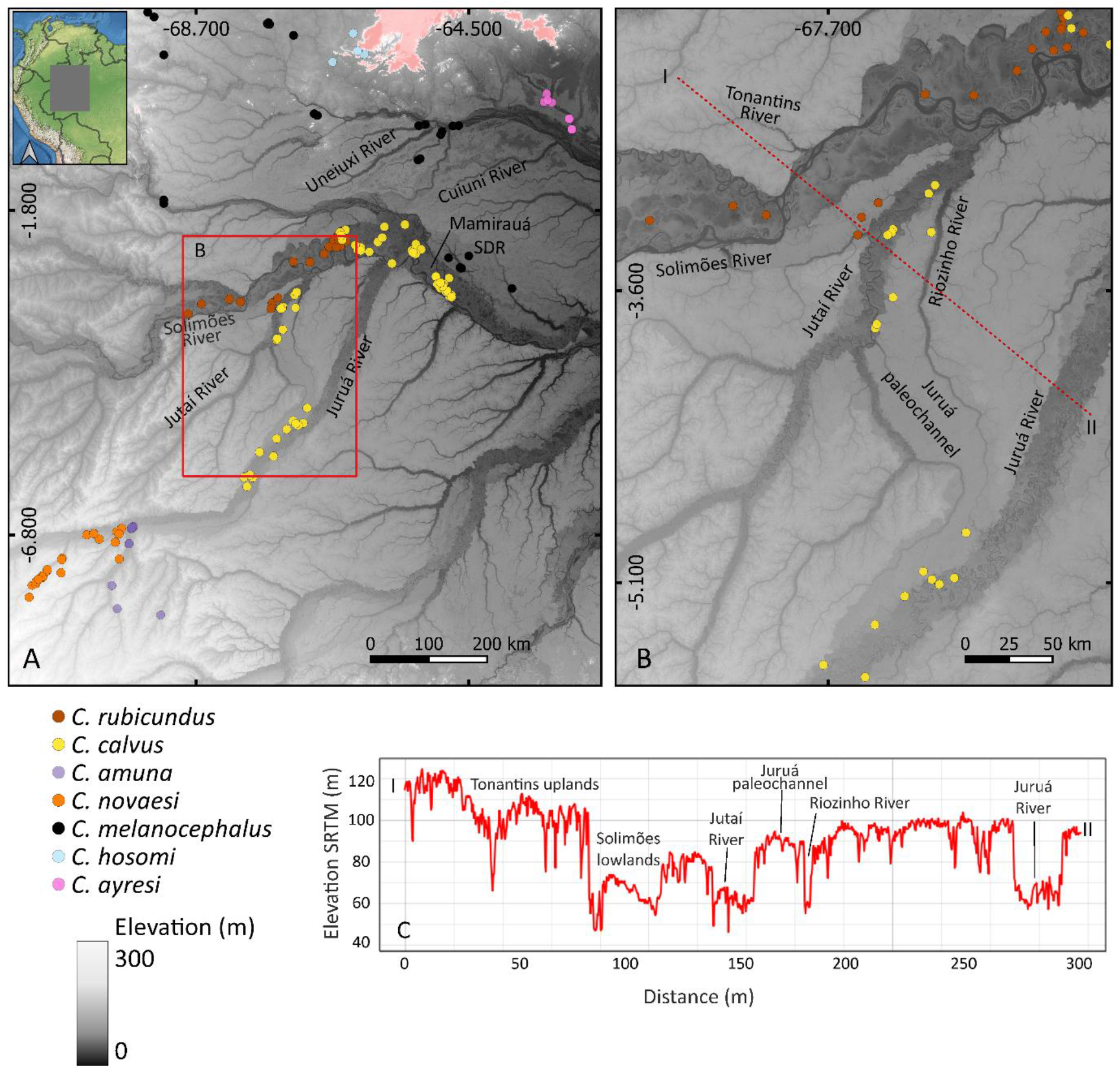
(A) The topography of the middle Solimões, Japurá, and Juruá River and the locality records of uakaris. (B) The influence of river rearrangements in the occurrence of *C. calvus* and *C. rubicundus*. Note that the geographic distribution of *C. calvus* overlaps the paleochannel connecting the middle Juruá River and the lower Jutaí River; and *C. rubicundus* occur in the lowlands of the Solimões River, with three disjunct populations occurring on opposite banks. The pointed red line in B is represented in the topographical profile.

We identified the presence of sharp lines that delimitate former floodplains with different elevations on both banks of the Solimões River (Figure 2). While a ∼80 km stretch of the Solimões River advanced in the NW direction towards the current position of the Tonantins River, it left behind alluvial deposits and a sequence of scroll bars, parallel streams (and narrow lakes), meanders, and oxbow lakes, forming a huge area of flooded forests on the southern bank (Figure 2). The occurrence of *C. rubicundus* overlaps with these floodplains and the disjunct distribution of this species can be explained by these intense tectonic and sedimentological activities.

*Cacajao calvus* and *C. rubicundus* occur in lowland areas below 100 m a.s.l., while *C. amuna* and *C. novaesi* occur both in seasonally flooded forests of the Tarauacá River and unflooded forests in areas > 200 m a.s.l. (Figure 2A). The separation of *C. calvus* and *C. rubicundus* from *C. amuna* and *C. novaesi* is consistent with geological and floristic discontinuities identified in the Juruá river basin (Higgins et al., 2011, Zuquim et al., 2021).

### Population genetic variation in *Cacajao*

In the PCA analysis, 61.5% of the genetic variation is explained by the first two components, with a clear distinction between black and bald-headed uakaris (Figure S2). When looking into each of these groups separately, we identified two clusters of *C. calvus* (populations from the Mamirauá SDR and the Jutaí River) and a cluster of *C. amuna*. In addition, *Cacajao rubicundus, C. ucayalii*, and *C. novaesi* individuals fell into three well-separated clusters, with the two first components explaining 32.9% of the genetic variation (Figure S3). We found four clusters of black uakaris, which include *C. hosomi, C. ayresi*, and the two populations of *C. melanocephalus* – the two first components explained 56.6% of the genetic variation (Figure S3). In the Japurá-Negro interfluve, a paleochannel – currently occupied by the Urubaxi and lower Cuiuni rivers – separates the two clusters found in *C. melanocephalus* (populations “melano” and “melanoW”; Figure 2C).

In the TreeMix analysis, we identified signals of shared ancestrality between *C. rubicundus* and *C. calvus*, especially involving the populations of both species from the Jutaí River. We also identified signals of shared ancestrality between *C. amuna* and *C. novaesi*, both occurring on opposite sides of the Tarauacá River (Figure 3). For the black uakaris, we identified signals of shared ancestrality between the populations of *C. melanocephalus* (melW) and *C. ayresi, C. melanocephalus* (mel) and *C. hosomi* (Figure 3).

**FIGURE 3.**
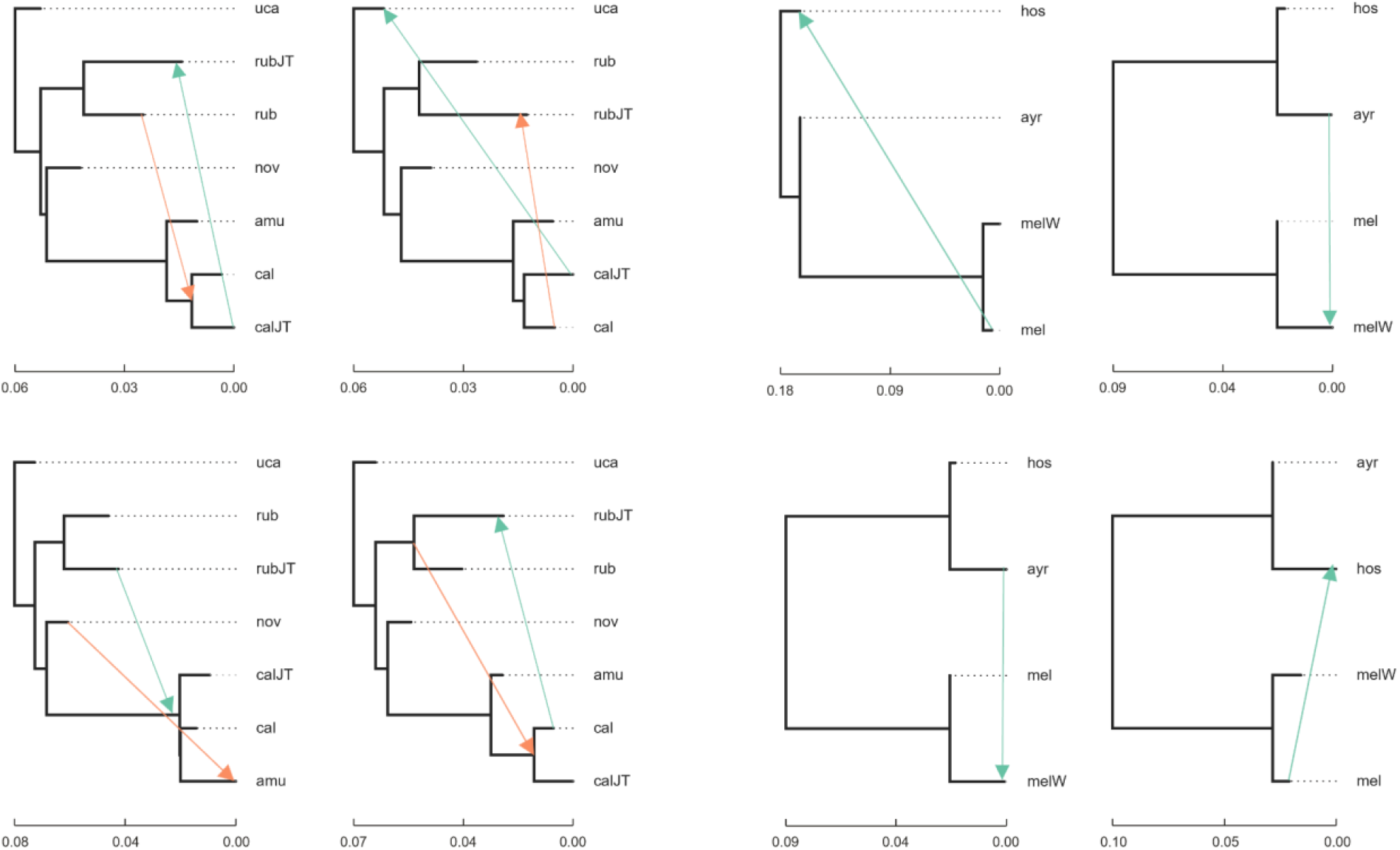
Shared ancestrality among the *Cacajao* species inferred in the TreeMix and based on the maximum likelihood tree. Gene flow events are depicted with arrows.

### Historical demography

Comparisons between the alternative demographic models suggest isolation with gene flow between *C. calvus* and *C. amuna*, but no gene flow between *C. calvus* and *C. rubicundus* and a population decline (Model 4) as the most likely to explain the observed data (Figure 4, Table S3). Parameter estimates based on the best-fit model suggest a large ancestral population size (NA1) of 438,282 (CI 95%: 487,932 – 712,027), with divergence times spanning the Middle Pleistocene (0.770 - 0.126 Ma), and first divergence (Tdiv1) between *C. rubicundus* and the ancestor of *C. calvus* and *C. amuna* beginning 763,009 (CI 95%: 440,471 – 1,204,923) years ago (Table S3). The second divergence event (Tdiv2), this time between *C. calvus* and *C. amuna*, occurred around 389,658 (CI 95%: 270,822 – 415,240) years ago. Model 4 also supports a scenario of exponential population decline for all three taxa, with times centred during the Late Pleistocene, between 37,575 and 68,077 years ago (Table 1). *Cacajao calvus* showed a higher rate of exponential population decline (GcalJT = 0.00173) compared to *C. rubicundus* (Grub = 0.00023) and *C. amuna* (Gamu = 0.00088), indicating a more dramatic population reduction. Current effective population size varied among taxa, with *C. rubicundus* being the largest with Ne 84,276, followed by *C. calvus* with Ne 35,094, and *C. amuna* with the smallest Ne values of 11,845. The migration rates per generation between *C. calvus* and *C. amuna* were relatively high, ranging between 1.12 (McalJT → Mamu) and 3.94 (McalJT ← Mamu). These migration rates were also considerably symmetrical, showing an overlap in the 95% confidence interval values (Table 1).

**TABLE 1.** Demographic parameters estimation values obtained by simulations based on the best-fit demographic model (Model 4) selected on FASTSIMCOAL.

| Parameters | Demographic units | Prior distribution | Estimates based on $\mu = 8.48e^{-9}$ (CI 95%) |
| --- | --- | --- | --- |
| Nrub | Current *C. rubicundus* population size | $10^4 - 10^5$ | 84,276 (24,880 – 88,091) |
| Narub | Ancestral *C. rubicundus* population size | $10^5 - 10^6$ | 356,747 (333,948 – 698,005) |
| Grub | *C. rubicundus* exponential population growth rate | $\log(\text{Nrub ratio})/\text{Tdiv}$ | 0.00023 (0.00003 – 0.00092) |
| NcalJT | Current *C. calvus* population size | $10^4 - 10^5$ | 35,094 (13,881 – 50,920) |
| NacalJT | Ancestral *C. calvus* population size | $10^5 - 10^6$ | 245,948 (228,292 – 309,057) |
| GcalJT | *C. calvus* exponential population growth | $\log(\text{NcalJT ratio})/\text{Tdiv}$ | 0.00173 (0.00021 – 0.00354) |
| Namu | Current *C. amuna* population size | $10^4 - 10^5$ | 11,845 (9,760 – 36,601) |
| Naamu | Ancestral *C. amuna* population size | $10^5 - 10^6$ | 257,911 (109,047 – 310,543) |
| Gamu | *C. amuna* exponential population growth rate | $\log(\text{Namu ratio})/\text{Tdiv}$ | 0.00088 (0.00073 – 0.00236) |
| NA1 (rub calJT, amu) | Ancestral size before first divergence time | $10^6 - 5 \times 10^6$ | 438,282 (487,932 – 712,027) |
| NA2 (calJT, amu) | Ancestral size before second divergence time | $5 \times 10^6 - 10^7$ | 884,027 (797,025 – 939,569) |
| Tgrub | Time of population decline of *C. rubicundus* (years) | $10^4 - 5 \times 10^4$ | 68,077 (29,664 – 92,111) |
| TgcalJT | Time of population decline of *C. calvus* (years) | $10^4 - 5 \times 10^4$ | 54,412 (48,702 – 102,031) |
| Tgamu | Time of population decline of *C. amuna* (years) | $10^4 - 5 \times 10^4$ | 37,575 (18,482 – 40,274) |
| Tdiv1 (rub calJT, amu) | First divergence time (years) | $5 \times 10^5 - 10^6$ | 763,009 (440,741 – 1,204,923) |
| Tdiv2 (calJT, amu) | Second divergence time (years) | $10^5 - 5 \times 10^5$ | 389,658 (270,822 – 415,240) |
| McalJT à Mamu | Migration from *C. calvus* to *C. amuna* | 0.1 – 10 | 1.12 (0.044 – 3.30) |
| McalJT β Mamu | Migration from *C. amuna* to *C. calvus* | 0.1 – 10 | 3.94 (0.821 – 4.69) |

**FIGURE 4.**
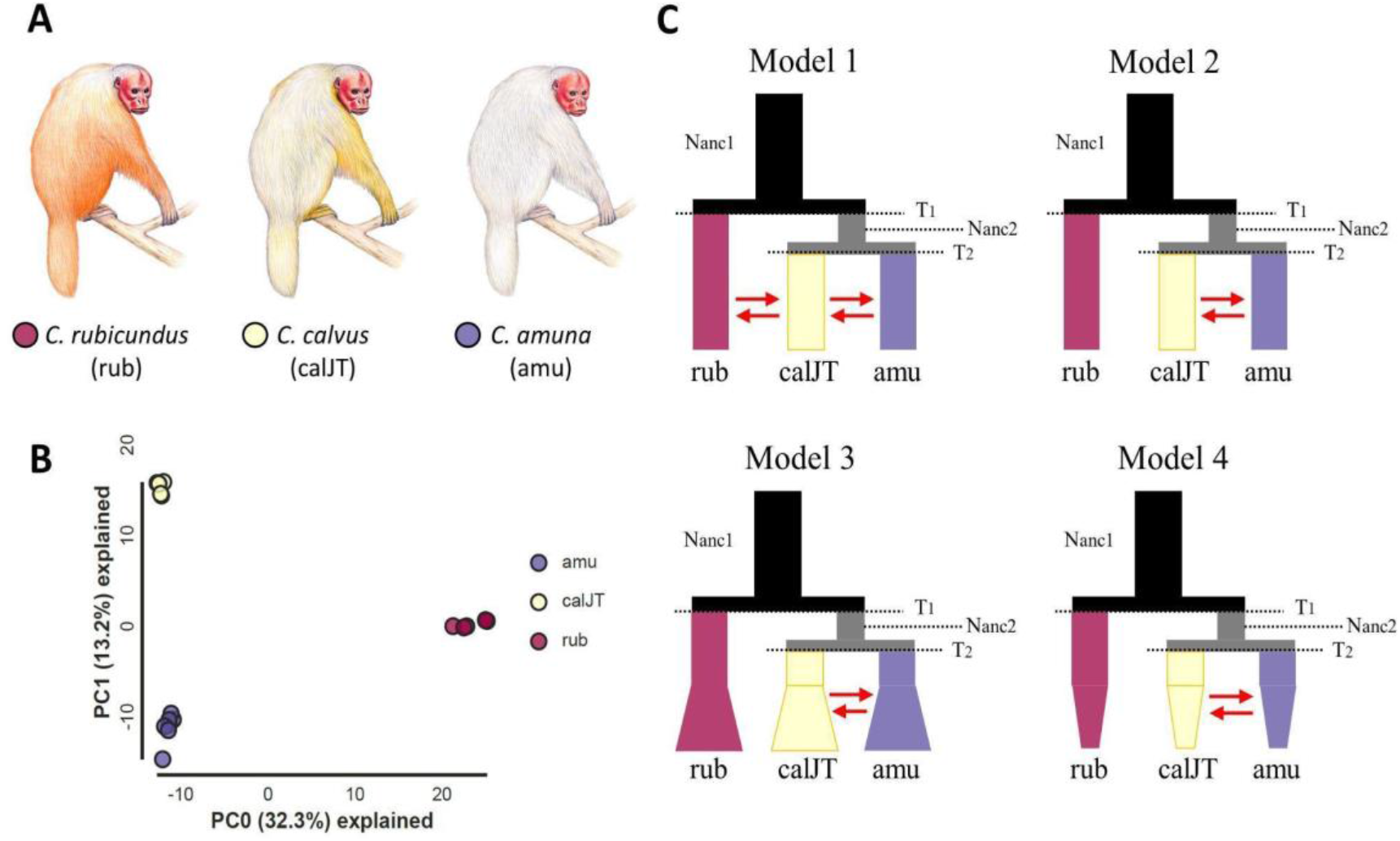
Population structure and historical demography scenarios were modelled among (A) *C. rubicundus* (rub), *C. calvus* (calJT), and *C. amuna* (amu). (B) PCA depicting allele variation among the three taxa and (C) the schematic representation of the four simulated demographic scenarios, where the best-fit model was Model 4 (see Table 2). Uakaris illustration by Stephen Nash.

## DISCUSSION

### The evolutionary history of *Cacajao*

We used genomics and landscape analyses to reconstruct the evolutionary history of the genus *Cacajao* and evaluate the role of landscape changes in western Amazonia on uakari populations. Our phylogeographic analysis indicated that the ancestral lineage of *Cacajao* was a lowland floodplain taxon that dispersed to different areas of western Amazonia in the mid-Pleistocene (from 1 Mya). By 0.7 Mya, the ancestral lineage of black uakaris occupied the Negro River basin to the north, while the bald-headed uakaris occupied the Jutaí River basin and extended to the middle Juruá River and west of the Peruvian forests about 0.4 Mya. Although we estimated a recent species-level divergence time (∼300 Kya), it predates landscape changes in the Solimões, Japurá and Juruá rivers (<100 Kya, Rukolainen et al., 2020; Rossetti et al., 2021). This scenario suggests that while the ancestral lineage dispersed through the Amazonian floodplains, passive disjunction caused by landscape and riverscape changes was essential in determining the current geographic distribution, population structure, and demographic history of the descendant lineages. A combination of dispersal ability (ecology) and earth history (landscape change) influenced *Cacajao* speciation at different timeframes. We posit, therefore, that the speciation in *Cacajao* was a product of the dispersal followed by vicariance, with the current diversity within the genus being a result of processes such as river rearrangements (Figure 5).

**FIGURE 5.**
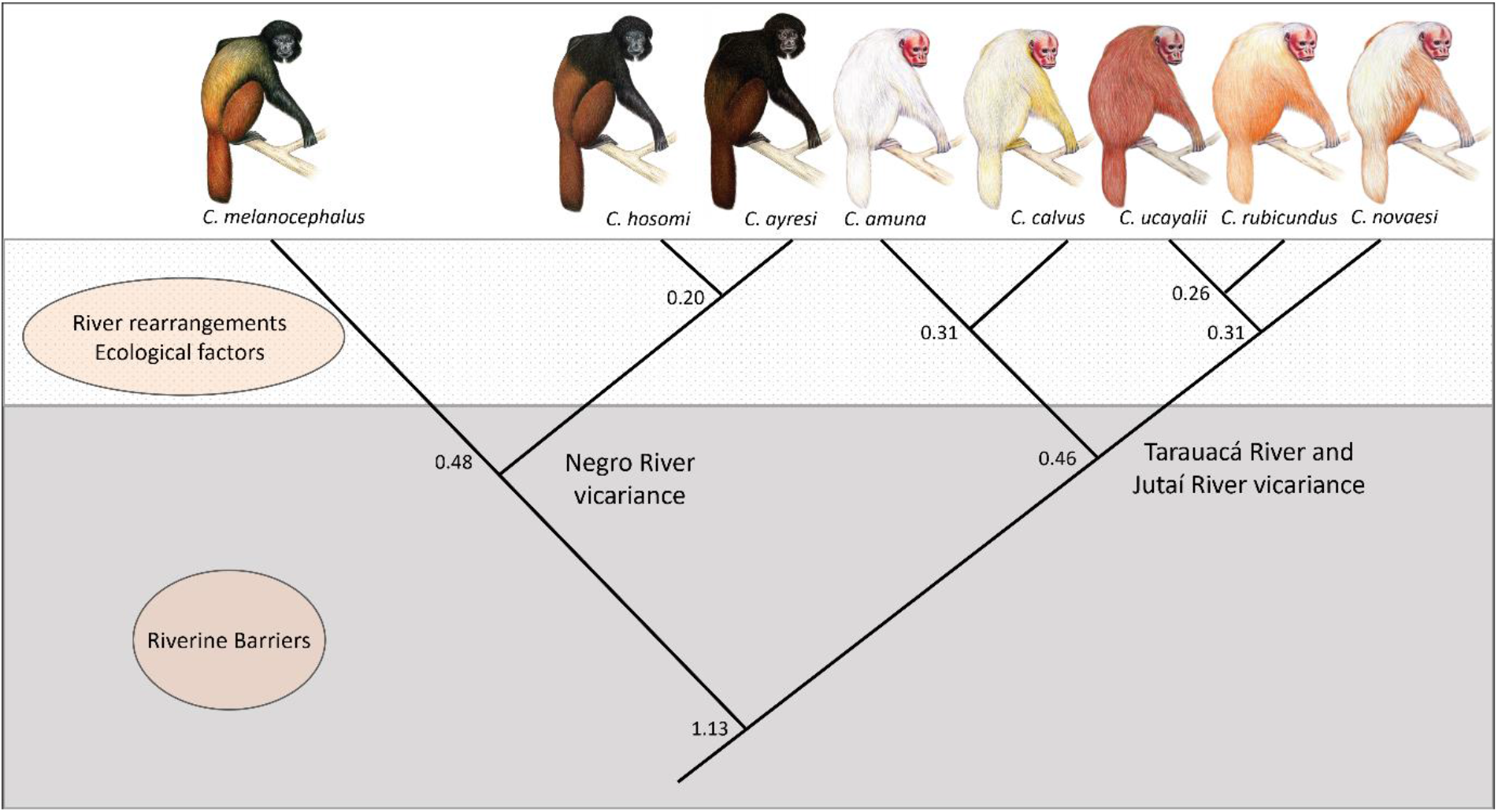
The main forces leading to the diversification in *Cacajao*. While the Negro, Tarauacá, and Jutaí rivers were important barriers to separating the ancestral lineages of black and bald-headed uakaris, processes like river rearrangements and ecological forces (e.g. retractions of the flooded forests, competition, and habitat requirements) were essential to maintain the separation of the recently diverged lineages and influenced their geographic distribution.

The Japurá River separated the two main *Cacajao* groups: black and bald-headed uakaris. Both lineages start to diverge at approximately 1 Mya. For black uakaris, the Negro River separated *C. melanocephalus* (South bank) and *C. hosomi* and *C. ayresi* (North bank), and their divergence start at approximately 0.48 Mya (Silva et al., 2022). The divergence of *Cacajao hosomi* and *C. ayresi*, however, is much more recent, 0.2 Mya (Silva et al., 2022), and competitive exclusion between bearded sakis and uakari monkeys has been considered a hypothesis for maintaining their separation (Boubli et al., 2008). *Cacajao* and *Chiropotes* are seed predators and their distribution is mostly allopatric (Ayres, 1989).

Changes in the landscape and riverscape of western Amazonia played a significant impact on the recently diverged lineages of bald-headed uakaris. Our analyses suggest that the ancestral lineage of *Cacajao amuna* arrived at the Tarauacá river basin at approximately 0.5 Mya. The dates of the landscape and riverscape changes in the Juruá River are uncertain, but recent studies suggest that the river flowed towards the Purus until approximately 0.1 Mya (Ruokolainen et al., 2020; Rossetti et al., 2021). This ancient Juruá River may have been responsible for speciation, with *C. amuna* and *C. calvus* diverging around 0.3 Mya due to geographic isolation on opposite sides of the river. In addition, this region is considered the boundary between the Miocene Pebas (Solimões) Formation, in the west, and the late Miocene–Pleistocene Içá Formation, to the east (Higgins et al., 2011), which is associated with the floristic turnovers zone identified in the Juruá river basin (Zuquim et al., 2021). However, the recent displacement of the Juruá River towards the Jutaí River basin (<0.1 Mya; see Ruokolainen et al., 2020; Rossetti et al., 2021) resulted in the isolation of *C. calvus* populations from the Jutaí River and the Mamirauá floodplains, and also may have promoted gene flow between *C. amuna* and *C. calvus*, as we identified in our analyses. Both species are entirely allopatric, separated by a distance of approximately 150 km (Silva et al., 2021, 2022). The population of *C. calvus* in the Jutaí River is now limited to a narrow floodplain area between the Jutaí-Riozinho interfluve (see Figure 2). The restricted distribution of this population is supported by several field surveys in the low Juruá River (e. g. Cardoso et al., 2014, Silva et al., 2021).

Tectonic activity and sedimentological processes promoted shifts in the Solimões River’s channel position during the Pleistocene-Holocene (Rossetti et al., 2021), potentially leading to the separation of the three populations of *C. rubicundus* that are entirely restricted to the flooded forests of this river. These shifts were responsible for the limited geographic distribution of the populations, with two on the North bank and one on the South bank. The region of the low Jutaí River was a meeting point for *C. rubicundus* and *C. calvus*, where occasional episodes of gene flow occurred due to facilitated contact between the two species caused by the landscape and riverscape changes. The restricted and disjunct geographic distribution of *C. calvus* and *C. rubicundus* populations and the retraction of the flooded forests (várzea) in western Amazonia during the late Pleistocene (see Pupim et al., 2019) led to episodes of population contractions in the last 70 Kya, as we found in our demographic analysis (Table 1). These events also explain the relatively low genetic diversity found in *Cacajao* (Kuderna et al., 2023).

### Ecological factors and genetic divergence in uakari monkeys

The bald-headed uakaris showed evidence of gene flow between *C. novaesi* and *C. amuna*, and between *C. rubicundus* and *C. calvus*. However, they are well-delimited in all analyses, including mitochondrial and ddRADseq, with a divergence time estimated at 0.46 Mya (Silva et al., 2022). Furthermore, they are separated by rivers that act as barriers to their dispersion, such as the Tarauacá River for *C. novaesi* and *C. amuna*, and the Jutaí River for *C. rubicundus* and *C. calvus* (Silva et al., 2021). In the case of *C. rubicundus* and *C. calvus*, a contact zone was reported in the Auati-Paraná channel (Vieira et al., 2008), with mixed groups occurring in a narrow area. The separation between these species can be maintained by the role of the Jutaí and Tarauacá rivers as barriers for their dispersal, as well as differences in niche requirements, potential interspecific competition, and hybridization.

The combination of landscape changes and ecological factors, such as retractions of the flooded forests, competition, and habitat characteristics, likely played a significant role in local adaptations in uakari monkeys. In this sense, the complex habitat affinity of uakari monkeys can generate opportunities for genetic divergence, especially when coupled with river rearrangements and the impact of this process on gene flow. This is evident from our gene flow results (Figures 3 and 4), which suggest that despite their divergence, there are signals of contact (or hybridization) among different taxa. The fact that these primates are not exclusive specialists of either the várzea or terra-firme forests is certainly observed in their complex pattern of diversification.

However, these factors remain largely uninvestigated in the *Cacajao* genus, which is still one of the least studied Neotropical primate genera. Long-term ecological studies were conducted only in two field sites for black uakaris: Jaú National Park (Barnett et al., 2005; Bezerra et al., 2010, 2011) and Pico da Neblina National Park (Boubli, 1999; Boubli and Tokuda, 2008), both protected areas in Brazil, and two studies on bald-headed uakaris in Mamirauá Reserve, Brazil (Ayres, 1986), and Lago Preto Conservation Concession, Peru (Bowler & Bodmer, 2009, 2011). Considering the complexity and the dynamic of changes in the western Amazonia that underlie the evolutionary history of *Cacajao*, these studies represent only a small fraction (although essential) of our understanding of the ecology and behaviour of the uakari monkeys.

Data on the ecology and behaviour of *Cacajao* in these different habitats are essential to understand the role of other important ecological processes (e.g., competitive exclusion) in maintaining the separation of recently diverged species, such as *C. amuna* and *C. calvus, C. rubicundus* and *C. ucayalii*, and *C. hosomi* and *C. ayresi*. In this context, the river rearrangements and the complex ecology of the uakari monkeys can have significant implications for the evolutionary history and diversification of these primates in western Amazonia.

Our findings highlight the significance of using diverse evidence to explore evolutionary history. Due to the intricate nature of the Amazon Rainforest, geological, ecological, and molecular data must be combined to investigate the time frame for speciation in different taxonomic groups and identify the adaptive mechanisms that led to the high diversity of this biome. By studying the effects of landscape and riverscape changes on the geographic distribution, population structure, and demographic history of the Amazonia biota, we can also gain insights into the species’ ability to adapt to future scenarios of anthropogenic changes.

## ACKNOWLEDGEMENTS

FES has received funding from the European Union’s Horizon 2020 research and innovation programme under the Marie Skłodowska-Curie grant agreement No 801505. FES also received funds from the Brazilian National Council for Scientific and Technological Development (CNPq) (Process nos.: 303286/2014-8, 303579/2014-5, 200502/2015-8, 302140/2020-4, 300365/2021-7, 301407/2021-5, 301925/2021-6); International Primatological Society (Conservation grant); The Rufford Foundation (14861-1, 23117-2, 38786-B), the Margot Marsh Biodiversity Foundation (SMA-CCO-G0023, SMA-CCOG0037), Primate Conservation Inc. (#1713 and #1689). SD acknowledges support from the *Fonds National de la Recherche Scientifique* (F.R.S.-FNRS, Belgium; grant n°F.4515.22), the Research Foundation - Flanders (*Fonds voor Wetenschappelijk Onderzoek* - Vlaanderen, FWO, Belgium; grant n°G098321N), and the European Union Horizon 2020 project MOOD (grant agreement n°874850). This research was also supported by the Gordon and Betty Moore Foundation (Grant #5344 to the Mamirauá Institute for Sustainable Development). JPB and RB were supported by NERC Standard Grant ‘Rise of the continent of the monkeys’ (NE/T000341/1).

## DATA AVAILABILITY STATEMENT

Raw ddRAD sequences: GenBank BioProject ID PRJNA830637

## REFERENCES

Akaike, H. (1974). A new look at the statistical model identification. IEEE Transactions on Automatic Control, 19, 716–723.

Andrews, S. (2018). FastQC: A quality control tool for high throughput sequence data. http://www.bioinformatics.babraham.ac.uk/projects/fastqc

Antonelli, A., Zizka, A., Carvalho, F.A., Scharn, R., Bacon, C.D., Silvestro, D., & Condamine, F.L. (2018). Amazonia is the primary source of Neotropical biodiversity Proceedings of the National Academy of Sciences USA, 115, 6034–6039.

Ayres, J.M. (1986). Uakaris and Amazonian Flooded Forest. PhD thesis, University of Cambridge, UK.

Ayres, J.M. (1989). Comparative feeding ecology of the uakari and bearded saki, Cacajao and Chiropotes. Journal of Human Evolution, 18, 697–716.

Ayres, J. M., & Clutton-Brock, T. H. (1992). River boundaries and species range size in Amazonian primates. American Naturalist, 140, 531–537.

Barbosa, W.E.S., Ferreira, M., Schultz, E.D., Luna, L.W., Larajeiras, T.O., Aleixo, A., & Ribas, C.C. (2022). Habitat association constrains population history in two sympatric ovenbirds along Amazonian floodplains. Journal of Biogeography, 49, 1683–1695.

Barnett, A.A., de Castilho, C.V., Shapley, R.L., & Anicácio, A. (2005). Diet, habitat selection and natural history of Cacajao melanocephalus ouakary in Jaú National Park, Brazil. International Journal of Primatology, 26, 949–969.

Barnett, A.A., Bowler, M., Bezerra, B.M., Defler, T.R. (2013). Ecology and behavior of uacaris (genus Cacajao). In L. Veiga, A.A. Barnett, S.F. Ferrari, M.A. Norconk (Eds.) Evolutionary biology and conservation of titis, sakis and uacaris (pp. 151–172). Cambridge, UK: Cambridge University Press.

Beck, R.M.D., de Vries, D., Janiak, M.C., Goodhead, I.B., & Boubli, J.P. (2023). Total evidence phylogeny of platyrrhine primates and a comparison of undated and tip-dating approaches. Journal of Human Evolution, 174: 103293.

Bezerra, B.M., Souto, A.S., & Jones, G. (2010). Responses of golden-backed uakaris, Cacajao melanocephalus, to call playback: implications for surveys in the flooded Igapó forest. Primates 51, 327–336.

Bezerra, B.M., Barnett, A.A., Souto, A., & Jones, G. (2011). Ethogram and natural history of golden-backed uakaris (Cacajao melanocephalus). International Journal of Primatology, 32, 46–68.

Boubli, J.P. (1999). Feeding ecology of black-headed uacaris (Cacajao melanocephalus melanocephalus) in Pico da Neblina National Park, Brazil. International Journal of Primatology, 20, 719–749.

Boubli, J.P., & Tokuda, M. (2008). Socioecology of black uakari monkeys, Cacajao hosomi, in Pico da Neblina National Park, Brazil. The role of the peculiar spatial–temporal distribution of resources in the Neblina forests. Primate Reports, 75, 3–10.

Boubli, J.P., da Silva, M.N.F., Amado, M.V., Hrbek, T., Pontual, F.B., & Farias, I.P. (2008). A taxonomic reassessment of Cacajao melanocephalus Humboldt (1811), with the description of two new species. International Journal of Primatology, 29, 723–741.

Boubli, J.P., Ribas, C., Alfaro, J.W.L., Alfaro, M.E., da Silva, M.N.F., Pinho, G.M., & Farias, I.P. (2015). Spatial and temporal patterns of diversification on the Amazon: a test of the riverine hypothesis afor all diurnal primates of Rio Negro and Rio Branco in Brazil. Molecular Phylogenetics and Evolution, 82, 400–412.

Bowler, M., & Bodmer, R. (2009). Social behavior in fission-fusion groups of red uakari monkeys (Cacajao calvus ucayalii). American Journal of Primatology, 71, 976–987.

Bowler, M., & Bodmer RE (2011). Diet and food choice in Peruvian red uakaris (Cacajao calvus ucayalii): selective or opportunistic seed predation? International Journal of Primatology, 32, 1109–1122.

Camargo, A., Werneck, F.P., Morando, M., Sites Jr., J.W., & Avila, L.J. (2013). Quaternary range and demographic expansion of Liolaemus darwinii (Squamata: Liolaemidae) in the Monte Desert of Central Argentina using Bayesian phylogeography and ecological niche modelling. Molecular Ecology, 22, 4038–4054.

Cardoso, N. A., Valsecchi, J., Vieira, T., & Queiroz, H.L. (2014). New records and range expansion of the white bald uakari (Cacajao calvus calvus, I. Geoffroy, 1847) in Central Brazilian Amazonia. Primates, 55, 199–206.

Catchen, J., Hohenlohe, P.A., Bassham, S., Amores, A., & Cresko, W.A. (2013). Stacks: an analysis tool set for population genomics. Molecular Ecology, 22, 3124–3140.

Catchen, J.M., Amores, A., Hohenlohe, P., Cresko, W., & Postlethwait, J.H. (2011). Stacks: building and genotyping loci de novo from short-read sequences. G3 Genes|Genomes|Genetics, 1, 171–182.

Cracraft, J., Ribas, C. C., d’Horta, F. M., Bates, J., Almeida, R. P., Aleixo, A.,…;Fritz, S. C. (2020). The origin and evolution of Amazonian species diversity. In V. Rull & A. Carnaval (Eds.), Neotropical diversification: patterns and processes. Fascinating life sciences (pp. 225–244). Cham: Springer Nature.

Dellicour, S., Rose, R., Faria, N. R., Lemey, P., & Pybus, G.O. (2016a). SERAPHIM: studying environmental rasters and phylogenetically informed movements, Bioinformatics, 32 (20), 3204–3206.

Dellicour, S., Rose, R. & Pybus, O.G. (2016b). Explaining the geographic spread of emerging epidemics: a framework for comparing viral phylogenies and environmental landscape data. BMC Bioinformatics, 17, 82.

Eaton, D.A.R., & Overcast, I. (2020). ipyrad: Interactive assembly and analysis of RADseq datasets. Bioinformatics, 36(8), 2592–2594.

Excoffier, L., Dupanloup, I., Huerta-Sanchez, E., Sousa, V. C., & Foll, M. (2013). Robust demographic inference from genomic and SNP data. PloS Genetics, 9, e1003905.

Fordham, G., Shanee, S., & Peck, M. (2020). Effect of river size on Amazonian primate community structure: a biogeographic analysis using updated taxonomic assessments. American Journal of Primatology, 82, e23136.

Gascon, C., Malcolm, J. R., Patton, J. L., da Silva, M. N., Bogart, J. P., Lougheed, S. C.,…Boag, P.T. (2000). Riverine barriers and the geographic distribution of Amazonian species. Proceedings of the National Academy of Sciences USA, 97, 13672–13677.

Gernhard, T. (2008). The conditioned reconstructed process. Journal of Theoretical Biology, 253, 769–778.

Gutenkunst, R. N., Hernandez, R. D., Williamson, S. H., & Bustamante, C. D. (2009). Inferring the joint demographic history of multiple populations from multidimensional SNP frequency data. PLoS Genetics, 5, 1–11.

Heymann, E.W., & Aquino, R. (2010). Peruvian red uakaris (Cacajao calvus ucayalii) are not flooded-forest specialists. International Journal of Primatology, 31, 751–758.

Higgins, M.A., Ruokolainen, K., Tuomisto, H., Llerena, N., Cardenas, G., Phillips, O.L., Vásquez, R., & Räsänen, M. (2011). Geological control of floristic composition in Amazonian forests. Journal of Biogeography, 38(11), 2136–2149.

Janiak, M. C., Silva, F. E., Beck, R. M. D., de Vries, D., Kuderna, L. F. K., Torosin, N. S.,…;Boubli, J. P. (2022). Two hundred and five newly assembled mitogenomes provide mixed evidence for rivers as drivers of speciation for Amazonian primates. Molecular Ecology, 31, 3888–3902.

Kalyaanamoorthy, S., Minh, B.Q., Wong, T.K.F., von Haeseler, A., & Jermiin, L.S. (2017). ModelFinder: Fast model selection for accurate phylogenetic estimates. Nature Methods, 14, 587–589.

Katoh, K., Rozewicki, J., & Yamada, K.D. (2019). MAFFT online service: multiple sequence alignment, interactive sequence choice and visualization, Briefings in Bioinformatics, 20(4), 1160–1166.

Kuderna, L.F., Gao, H., Janiak, M.C., Kuhlwilm, M., Orkin, J.D., Bataillon, T., Manu, S., Valenzuela, A., Bergman, J., Rouselle, M.,…Silva, F.E. (2023). A global catalog of whole-genome diversity from 233 primate species. bioRxiv, 1–5.

Lemey, P., Rambaut, A., Welch, J.J., & Suchard, M.A. (2010). Phylogeography takes a relaxed random walk in continuous space and time. Molecular Biology and Evolution, 27, 1877–1885.

Luna, L.W., Ribas, C.C., & Aleixo, A. (2022). Genomic differentiation with gene flow in a widespread Amazonian floodplain-specialist bird species. Journal of Biogeography, 49(9), 1670–1682.

Luna, L.W., Naka, L.N., Thom, G., Knowles, L.L., Sawakuchi, A.O., Aleixo, A. & Ribas, C.C., (2023). Late Pleistocene landscape changes and habitat specialization as promoters of population genomic divergence in Amazonian floodplain birds. Molecular Ecology, 32(1), 214–228.

Lynch-Alfaro, J.W., Boubli, J.P., Paim, F.P., Ribas, C.C., da Silva, M.N.F., Messia, M.R.,…Farias, I.P. (2015). Biogeography of squirrel monkeys (genus Saimiri): South-Central Amazon origin and rapid pan-Amazonian diversification of a lowland primate. Molecular Phylogenetics and Evolution, 82, 436–454.

McHugh, S., Cornejo, F., McKibben, J., Zarate, M., Tello, C., Jiménez, C., & Schmitt, C. (2020). First record of the Peruvian yellow-tailed woolly monkey Lagothrix flavicauda in the Región Junín, Peru. Oryx, 54(6), 814–818.

Miranda, C.L., Farias, I.P., Da Silva, M.N.F., Antonelli, A., Machado, A.F., Leite, R.N.,…;Pieczarka, J.C., (2022). Diversification of Amazonian spiny tree rats in genus Makalata (Rodentia, Echimyidae): cryptic diversity, geographic structure and drivers of speciation. PLoS One, 17(12), e0276475.

Mourthé, Í., Hilário, R.R., Carvalho, W.D., & Boubli JP (2022). Filtering effect of large rivers on primate distribution in the Brazilian Amazonia. Frontiers in Ecology and Evolution, 10, 857920.

Nascimento, F.F., Lazar, A., Menezes, A.N., Durans, A.D.M., Moreira, J.C., Salazar-Bravo, J., D′ Andrea, P.S.,…Bonvicino, C.R. (2013). The role of historical barriers in the diversification processes in open vegetation formations during the Miocene/Pliocene using an ancient rodent lineage as a model. PLoS One, 8, e61924.

Oberdorff, T., Dias, M.S., Jézéquel, C., Albert, J.S., Arantes, C.C., Bigorne, R.,…Zuanon, J. (2019). Unexpected fish diversity gradients in the Amazon basin. Science Advances, 11, 5(9), eaav8681.

Peterson, B.K., Weber, J.N., Kay, E.H., Fisher, H.S., & Hoekstra, H.E. (2012). Double digest RADseq: an inexpensive method for de novo SNP discovery and genotyping in model and non-model species. PLoS One, 7, 1–11.

Pickrell, J., & Pritchard, J. (2012). Inference of population splits and mixtures from genome-wide allele frequency data. Nature Precedings, 1–1.

Pupim, F.N., Sawakuchi, A.O., Almeida, R.P.D., Ribas, C.C., Kern, A.K., Hartmann, G.A,.…Grohmann, C.H. (2019). Chronology of Terra Firme formation in Amazonian lowlands reveals a dynamic Quaternary landscape. Quaternary Science Reviews, 210, 154–163.

Pybus, O.G., Suchard, M.A., Lemey, P., Bernardin, F.J., Rambaut, A., Crawford, F.W.,…Delwart, E.L. (2012). Unifying the spatial epidemiology and molecular evolution of emerging epidemics. Proceedings of the National Academy of Sciences USA, 109, 15066–15071.

QGIS Development Team (2021). QGIS Geographical Information System. Open Source. Geospatial Foundation Project.

Rambaut, A., Drummond, A.J., Xie, D., Baele, G., & Suchard, M.A. (2018). Posterior summarization in Bayesian phylogenetics using Tracer 1.7. Systematic Biology, 67, 901–904.

Rochette, N.C., & Catchen, J.M. (2017). Deriving genotypes from RAD-seq short-read data using Stacks. Nature Protocols, 12, 2640–2659.

Rossetti, D.F. (2014). The role of tectonics in the late Quaternary evolution of Brazil’s Amazonian landscape. Earth-Science Reviews, 139, 362–389.

Rossetti, D.F., Vasconcelos D.L., Valeriano M.M., & Bezerra F.H.R. (2021). Tectonics and drainage development in Central Amazonia: the Juruá River. Catena, 206, 105560.

Ruokolainen, K., Moulatlet, G.M., Zuquim, G., Hoorn, C., & Tuomisto, H. (2020). Geologically recent rearrangements in central Amazonian river network and their importance for the riverine barrier hypothesis. Frontiers in Biogeography, 11(3), e45046.

Sawakuchi, A.O., Schultz, E.D., Pupim, F.N., Bertassoli Jr, D.J., Souza, D.F., Cunha, D.F.,…Ribas, C.C. (2022). Rainfall and sea level drove the expansion of seasonally flooded habitats and associated bird populations across Amazonian. Nature Communications, 13, 4945.

Silva, S.M., Peterson, A.T., Carneiro, L., Burlamaqui, T.C.T., Ribas, C.C., Sousa-Neves, T.,…Batista, R., (2019). A dynamic continental moisture gradient drove Amazonian bird diversification. Science Advances, 5(7), eaat5752.

Silva, F.E., Lemos, L.P., Ravetta, A.L., Röhe, F., Sampaio, R., Franco, C.L.P.,…Boubli, J.P., (2021). On the geographic distribution of the bald uakaris (Cacajao calvus ssp.) in Brazilian Amazonia. Primate Conservation (35), 69–86.

Silva, F.E., Valsecchi, J., Roos, C., Bowler, M., Röhe, F., Sampaio, R.,…Boubli, J. (2022). Molecular phylogeny and systematics of bald uakaris, genus Cacajao Lesson, 1840 (Primates: Pitheciidae), with the description of a new species. Molecular Phylogenetics and Evolution, 173, 107509.

Silva-Júnior, J.S., Figueiredo-Ready, W.M.B., & Ferrari, S.F.S. (2013). Taxonomy and geographic distribution of the Pithecidae. In L. Veiga, A.A. Barnett, S.F. Ferrari, M.A. Norconk (Eds.) Evolutionary biology and conservation of titis, sakis and uacaris (pp. 31–42). Cambridge, UK: Cambridge University Press.

Suchard, M.A., Lemey, P., Baele, G., Ayres, D.L., Drummond, A.J., & Rambaut, A. (2018). Bayesian phylogenetic and phylodynamic data integration using BEAST 1.10. Virus Evolution, 4, 1–5.

Thom, G., Xue, A.T., Sawakuchi, A.O., Ribas, C.C., Hickerson, M.J., Aleixo, A., & Miyaki, C., 2020. Quaternary climate changes as speciation drivers in the Amazon floodplains. Science Advances, 6(11), p.eaax4718.

Vermeer, J., Tello-Alvarado, J.C., del Castillo, J.T.V., & Bóveda-Penalba, A.J. (2013). A new population of red uakaris (Cacajao calvus spp.) in the mountains of North-Eastern Peru. Neotropical Primates, 20, 12–17.

Vieira, T., Oliveira, M., & Queiroz, H. (2008). Novas informações sobre a distribuição de Cacajao calvus na Reserva de Desenvolvimento Sustentável Mamirauá. Uakari, 4, 41–51.

Wallace, A. R. (1852). On the monkeys of the Amazon. Proceedings of the Zoological Society of London, 20, 107–110.

Werneck, F.P., Leite, R.N., Geurgas, S.R., & Rodrigues, M.T. (2015). Biogeographic history and cryptic diversity of saxicolous Tropiduridae lizards endemic to the semiarid Caatinga. BMC Evolutionary Biology, 15, 1–24.

Zuquim, G., Tuomisto, H., Chaves, P.P., Emilio, T., Moulatlet, G.M., Ruokolainen, K.,…Balslev, H. (2021). Revealing floristic variation and map uncertainties for different plant groups in western Amazonia. Journal of Vegetation Science, 32:e13081.

