## Supplementary Material for "The impact of Quaternary Amazonian river dynamics on patterns and process of diversification in uakari monkeys (genus *Cacajao*)"

**TABLE S1. Samples used in the molecular analyses and their origin.** Collection codes refer to the following zoological collections: Museu Paraense Emílio Goeldi (MPEG), Instituto de Desenvolvimento Sustentável Mamirauá (IDSM), Instituto Nacional de Pesquisas da Amazônia (INPA), Museu de Zoologia da Universidade de São Paulo (MUZUSP), Field Museum of Natural History (FMNH), Museum of Vertebrate Zoology at UC-Berkeley (MVZ), Universidade Federal do Pará (UFPA), Universidade Federal do Amazonas - Laboratório de Evolução e Genética Animal (CTGAM).

| **Tree code** | **Taxon** | **Cyt *b*** | **ddRAD** | **Collection code** | **Field code** | **Gene Bank code (nucleotide/SRA)** | **Locality** | **Long** | **Lat** | **Reference** |
|---|---|---|---|---|---|---|---|---|---|---|
| Cacajao_sp_IDSM03668_Tar | *C. amuna* | ✓ | ✓ | IDSM03668 | FES64 | ON337703/ SAMN27734981 | Tarauacá River, right bank, Brazil | -69.925 | -6.753 | Silva et al. 2022 |
| Cacajao_sp_IDSM03674_Tar | *C. amuna* | ✓ | ✓ | IDSM03674 | FES70 | ON337704/ SAMN27734982 | Tarauacá River, right bank, Brazil | -69.667 | -6.671 | Silva et al. 2022 |
| Cacajao_sp_IDSM03675_Tar | *C. amuna* | ✓ | ✓ | IDSM03675 | FES71 | ON337705/ SAMN27734983 | Tarauacá River, right bank, Brazil | -69.667 | -6.671 | Silva et al. 2022 |
| Cacajao_sp_IDSM03676_Tar | *C. amuna* | ✓ | ✓ | IDSM03676 | FES95 | ON337706/ SAMN27734984 | Itucumã Lake, Tarauacá River, right bank, Brazil | -69.738 | -6.935 | Silva et al. 2022 |
| Cacajao_sp_INPA007279_Pauini | *C. amuna* | ✓ | ✓ | INPA007279 | RS62 | ON337709/ SAMN27734986 | Pauini River, left bank, Brazil | -69.132 | -7.605 | Sampaio et al. (2018); Silva et al. 2022 |
| Cacajao_sp_INPA007276_Pauini | *C. amuna* | ✓ | ✓ | INPA007276 | RS63 | ON337708/ SAMN27734985 | Pauini River, left bank, Brazil | -69.132 | -7.605 | Sampaio et al. (2018); Silva et al. 2022 |
| Cacajao_sp_INPA007280_Pauini | *C. amuna* | ✓ | ✓ | INPA007280 | RS64 | ON337710/ SAMN27734987 | Pauini River, left bank, Brazil | -69.132 | -7.605 | Sampaio et al. (2018); Silva et al. 2022 |
| Cacajao_sp_INPA05241_Tar | *C. amuna* | ✓ | | INPA05241 | CCM112 | ON337707 | Tarauacá River, right bank, Brazil | -71.360 | -8.830 | Silva et al. 2022 |
| Cacajao_sp_MPEG21861_Jurupari | *C. amuna* | ✓ | | MPEG21861 | | ON337711 | Jurupari River, Brazil | -70.158 | -7.579 | Silva Júnior and Martins (1999) |
| Cacajao_sp_MPEG21862_Jurupari | *C. amuna* | ✓ | | MPEG21862 | | ON337712 | Jurupari River, Brazil | -70.158 | -7.579 | Silva Júnior and Martins (1999) |
| C_calvus_IDSM00519_RDSM | *C. calvus* | ✓ | ✓ | IDSM00519 | Masto283 | ON337686/ SAMN27734995 | Mamirauá Reserve, Brazil | -64.935 | -2.912 | Silva et al. 2022 |
| C_calvus_IDSM00040_Jut | *C. calvus* | ✓ | ✓ | IDSM00040 | JT22 | ON337685/ SAMN27734994 | Jutaí River, right bank, Brazil | -67.395 | -3.313 | Silva et al. 2022 |
| C_calvus_IDSM00003_Jut | *C. calvus* | ✓ | ✓ | IDSM00003 | JT03 | ON337684/ SAMN27734993 | Jutaí River, right bank, Brazil | -67.374 | -3.300 | Silva et al. 2022 |
| C_calvus_IDSM00784_Jut | *C. calvus* | | ✓ | IDSM00784 | JT82 | SAMN27734996 | Jutaí River, right bank, Brazil | -67.15 | -3.06 | Silva et al. 2022 |
| C_calvus_IDSM00785_Jut | *C. calvus* | ✓ | ✓ | IDSM00785 | JT85 | ON337688/ SAMN27734997 | Riozinho, left bank, Brazil | -67.137 | -3.298 | Silva et al. 2022 |
| C_calvus_IDSM00786_Jut | *C. calvus* | | ✓ | IDSM00786 | JT88 | SAMN27734998 | Jutaí River, right bank, | -67.46 | -3.79 | Silva et al. 2022 |

| | | | | | | | | | | |
|---|---|---|---|---|---|---|---|---|---|---|
| | | | | | | | Brazil | | | |
| C_calvus_IDSM00787_Jut | *C. calvus* | | ✓ | IDSM00787 | JT90 | SAMN27734999 | Jutaí River, right bank, Brazil | -67.45 | -3.771 | Silva et al. 2022 |
| C_calvus_IDSM03677_RDSM | *C. calvus* | ✓ | ✓ | IDSM03677 | FES102 | ON337687/ SAMN27735001 | Mamirauá Reserve, Brazil | -64.854 | -3.071 | Silva et al. 2022 |
| C_calvus_IDSM03174_RDSM | *C. calvus* | | ✓ | IDSM03174 | Masto1384 | SAMN27735000 | Mamirauá Reserve, Brazil | -65.333 | -2.411 | Silva et al. 2022 |
| C_calvus_UFPA-Ccn1_Carauari | *C. calvus* | ✓ | | UFPA-Ccn1 | | FJ531666 | Carauari, Rio Juruá, Brazil | -66.963 | -4.974 | Figueiredo-Ready et al. 2013 |
| C_calvus_MZUSP17537_RDSM | *C. calvus* | ✓ | | MZUSP17537 | | FJ531651.1 | Mamirauá Reserve, Brazil | -64.935 | -3.000 | Figueiredo-Ready et al. 2013 |
| C_rub_IDSM00083_Jut | *C. rubicundus* | ✓ | ✓ | IDSM00083 | JT63 | ON337690/ SAMN27735020 | ESEC_Jutaí-Solimões, Brazil | -67.423 | -3.201 | Silva et al. 2022 |
| C_rub_IDSM00082_Jut | *C. rubicundus* | ✓ | ✓ | IDSM00082 | JT62 | ON337689/ SAMN27735019 | ESEC_Jutaí-Solimões, Brazil | -67.423 | -3.201 | Silva et al. 2022 |
| C_rub_IDSM00788_Jut | *C. rubicundus* | ✓ | ✓ | IDSM00788 | JT78 | ON337691/ SAMN27735021 | ESEC_Jutaí-Solimões, Brazil | -67.548 | -3.312 | Silva et al. 2022 |
| C_rub_IDSM03665_Ica | *C. rubicundus* | ✓ | ✓ | IDSM03665 | FES46 | ON337692/ SAMN27735022 | Jacurapá_River, left bank, Brazil | -68.618 | -3.237 | Silva et al. 2022 |
| C_rub_IDSM03666_Ica | *C. rubicundus* | ✓ | ✓ | IDSM03666 | FES47 | ON337693/ SAMN27735023 | Jacurapá_River, left bank, Brazil | -68.618 | -3.237 | Silva et al. 2022 |
| C_rub_IDSM03667_Ica | *C. rubicundus* | ✓ | ✓ | IDSM03667 | FES48 | ON337694/ SAMN27735024 | Jacurapá_River, left bank, Brazil | -68.618 | -3.237 | Silva et al. 2022 |
| C_rub_FJ531652_AP | *C. rubicundus* | ✓ | | MZUSP17552 | | FJ531652 | Buiuçu, Auatí-Paraná channel, Brazil | -66.447 | -2.353 | Figueiredo-Ready et al. 2013 |
| C_rub_FJ531653_AP | *C. rubicundus* | ✓ | | MZUSP17553 | | FJ531653 | Buiuçu, Auatí-Paraná channel, Brazil | -66.447 | -2.353 | Figueiredo-Ready et al. 2013 |
| C_ucayalii_IDSM03678_PNSD | *C. ucayalii* | ✓ | ✓ | IDSM03678 | FES100 | ON337695/ SAMN27735025 | Moa River, Serra do Divisor National Park (SDNP), Brazil | -73.668 | -7.461 | Silva et al. 2022 |
| C_ucayalii_IDSM03679_PNSD | *C. ucayalii* | ✓ | ✓ | IDSM03679 | FES101 | ON337696/ SAMN27735026 | Moa River, Serra do Divisor National Park (SDNP), Brazil | -73.668 | -7.461 | Silva et al. 2022 |
| C_ucayalii_FJ531660_EstEq | *C. ucayalii* | ✓ | | MPEG1848 | | FJ531660 | Estirão do Equador, Javari River, Brazil | -71.676 | -4.436 | Figueiredo-Ready et al. 2013 |
| C_ucayalii_FJ531662_EstEq | *C. ucayalii* | ✓ | | MPEG1849 | | FJ531662 | Estirão do Equador, Javari River, Brazil | -71.676 | -4.436 | Figueiredo-Ready et al. 2013 |
| C_ucayalii_FJ531663_Tapiche | *C. ucayalii* | ✓ | | UFPA-Ccu4957 | | FJ531663 | Tapiche River, Peru | -74.004 | -5.655 | Figueiredo-Ready et al. 2013 |
| C_ucayalii_MB1_Peru | *C. ucayalii* | ✓ | | | MB1 | ON337698 | Lago Preto | -71.765 | -4.458 | Silva et al. 2022 |

| | | | | | | | | | | |
|---|---|---|---|---|---|---|---|---|---|---|
| | | | | | | | Conservation Concession, Loreto, Peru | | | |
| C_ucayalii_MB12B_Peru | *C. ucayalii* | √ | | | MB12B | ON337697 | Lago Preto Conservation Concession, Loreto, Peru | -71.765 | -4.458 | Silva et al. 2022 |
| C_ucayalii_MB4_Peru | *C. ucayalii* | √ | | | MB4 | ON337700 | Lago Preto Conservation Concession, Loreto, Peru | -71.765 | -4.458 | Silva et al. 2022 |
| C_ucayalii_MB49_Peru | *C. ucayalii* | √ | | | MB49 | ON337699 | Lago Preto Conservation Concession, Loreto, Peru | -71.765 | -4.458 | Silva et al. 2022 |
| C_ucayalii_MB54_Peru | *C. ucayalii* | √ | | | MB54 | ON337701 | Lago Preto Conservation Concession, Loreto, Peru | -71.765 | -4.458 | Silva et al. 2022 |
| C_ucayalii_MB8_Peru | *C. ucayalii* | √ | | | MB8 | ON337702 | Lago Preto Conservation Concession, Loreto, Peru | -71.765 | -4.458 | Silva et al. 2022 |
| C_ucayalli_FJ531654_Tapiche | *C. ucayalii* | √ | | UFPA-Ccu4958 | | FJ531654 | Tapiche River, Peru | -74.004 | -5.655 | Figueiredo-Ready et al. 2013 |
| C_ucayalli_FJ531661_EstEq | *C. ucayalii* | √ | | MPEG1850 | | FJ531661 | Estirão do Equador, Javari River, Brazil | -71.676 | -4.436 | Figueiredo-Ready et al. 2013 |
| C_ucayalli_FJ531664_Tapiche | *C. ucayalii* | √ | | UFPA-Ccu4959 | | FJ531664 | Tapiche River, Peru | -74.004 | -5.655 | Figueiredo-Ready et al. 2013 |
| C_novaesi_IDSM03669_Eiru | *C. novaesi* | √ | √ | IDSM03669 | FES65 | ON375824/SAMN 27735014 | Igarapé Preto, Juruá River, right bank, Brazil | -64.800 | -3.117 | Silva et al. 2022 |
| C_novaesi_IDSM03670_Eiru | *C. novaesi* | √ | √ | IDSM03670 | FES66 | ON375825/SAMN 27735015 | Igarapé Preto, Juruá River, right bank, Brazil | -70.196 | -6.864 | Silva et al. 2022 |
| C_novaesi_IDSM03671_Eiru | *C. novaesi* | √ | √ | IDSM03671 | FES67 | ON375826/SAMN 27735016 | Igarapé Preto, Juruá River, right bank, Brazil | -70.196 | -6.864 | Silva et al. 2022 |
| C_novaesi_IDSM03672_Eiru | *C. novaesi* | √ | √ | IDSM03672 | FES68 | ON375827/SAMN 27735017 | Igarapé Preto, Juruá River, right bank, Brazil | -70.196 | -6.864 | Silva et al. 2022 |
| C_novaesi_IDSM03673_Eiru | *C. novaesi* | √ | √ | IDSM03673 | FES69 | ON375828/SAMN 27735018 | Igarapé Preto, Juruá River, right bank, Brazil | -69.925 | -6.753 | Silva et al. 2022 |
| C_ayresi_CTGAM5666_Araca | *C. ayresi* | √ | √ | CTGAM5666 | | ON337682/ SAMN27734988 | Acará River, left bank, Brazil | -62.950 | -0.380 | Bertuol 2015 |

| | | | | | | | | | | |
|---|---|---|---|---|---|---|---|---|---|---|
| C_ayresi_CTGAM5667_Araca | *C. ayresi* | ✓ | ✓ | CTGAM5667 | | ON337683/ SAMN27734989 | Acará River, left bank, Brazil | -62.950 | -0.380 | Bertuol 2015 |
| C_ayresi_INPA5246_Madixi | *C. ayresi* | ✓ | ✓ | INPA5246/ CTGAM5708 | JPB135 | EU560409.1/ SAMN27734990 | Igarapé Madixi, Brazil | -63.340 | -0.120 | Boubli et al. 2008 |
| C_ayresi_INPA5247_Araca | *C. ayresi* | ✓ | ✓ | INPA5247/ CTGAM5717 | JPB138 | EU560410.1/ SAMN27734991 | Acará River, left bank, Brazil | -62.910 | -0.540 | Boubli et al. 2008 |
| C_ayresi_INPA5248_Araca | *C. ayresi* | ✓ | ✓ | INPA5248/ CTGAM5721 | JPB139 | EU560411.1/ SAMN27734992 | Acará River, left bank, Brazil | -62.910 | -0.540 | Boubli et al. 2008 |
| C_hosomi_CTGAM5698_Imeri | *C. hosomi* | ✓ | ✓ | CTGAM5698 | JPB102 | EU560412/ SAMN27735002 | Serra do Imeri, Xamata, Brazil | -65.270 | 0.490 | Boubli et al. 2008 |
| C_hosomi_INPA5242_SGC | *C. hosomi* | ✓ | | INPA5242 | JPB001 | EU560418.1 | São Gabriel da Cacheira, Brazil | -66.110 | 0.610 | Boubli et al. 2008 |
| C_hosomi_INPA5249_Waputar | *C. hosomi* | ✓ | | INPA5249 | | EU560414.1 | Serra do Padre e Waputar, Brazil | -66.210 | 0.660 | Boubli et al. 2008 |
| C_hosomi_INPA5250_Waputar | *C. hosomi* | ✓ | | INPA5250 | JPB154 | EU560415.1 | Serra do Padre e Waputar, Brazil | -66.210 | 0.660 | Boubli et al. 2008 |
| C_hosomi_INPA5251_Waputar | *C. hosomi* | ✓ | | INPA5251 | JPB153 | EU560416.1 | Serra do Padre e Waputar, Brazil | -66.210 | 0.660 | Boubli et al. 2008 |
| C_hosomi_INPA5252_Waputar | *C. hosomi* | ✓ | | INPA5252 | JPB152 | EU560417.1 | Serra do Padre e Waputar, Brazil | -66.600 | 0.490 | Boubli et al. 2008 |
| C_hosomi_JPB163_Venez | *C. hosomi* | ✓ | | | JPB163 | ON375823 | Venezuela | -65.280 | 2.250 | Bertuol 2015 |
| C_hosomi_CTGAM5716_Imeri | *C. hosomi* | ✓ | ✓ | CTGA5716 | JPB102 | EU560413/ SAMN27735002 | Serra do Imeri, Xamata, Brazil | -65.270 | 0.490 | Boubli et al. 2008 |
| C_melano_CTGAM5663_R.Negro | *C. melanocephalus* | ✓ | ✓ | CTGAM5663 | | ON337675/ SAMN27735005 | Negro River, right bank, Brazil | -64.740 | -0.490 | unpubl. data |
| C_melano_CTGAM5665_R.Negro | *C. melanocephalus* | ✓ | ✓ | CTGAM5665 | | ON337676/ SAMN27735006 | Negro River, right bank, Brazil | -64.650 | -0.490 | unpubl. data |
| C_melano_CTGAM65_R.Negro | *C. melanocephalus* | ✓ | ✓ | CTGAM0065 | SIS65 | ON337677/ SAMN27735010 | Igarapé Parati, Negro River, right bank, Brazil | -64.910 | -0.580 | unpubl. data |
| C_melano_CTGAM756_Jap | *C. melanocephalus* | ✓ | ✓ | CTGAM0756 | | ON337678/ SAMN27735004 | Japurá River, left bank, Brazil | -69.200 | -1.690 | unpubl. data |
| C_melano_CTGAM757_Jap | *C. melanocephalus* | ✓ | ✓ | CTGAM0757 | JAP757 | ON337679/ SAMN27735011 | Japurá River, left bank, Brazil | -69.200 | -1.690 | unpubl. data |
| C_melano_CTGAM775_Jap | *C. melanocephalus* | ✓ | ✓ | CTGAM0775 | JAP775 | ON337680/ SAMN27735012 | Japurá River, left bank, Brazil | -69.340 | -1.660 | unpubl. data |
| C_melano_CTGAM98_Aiuana | *C. melanocephalus* | ✓ | ✓ | CTGAM0098 | SIS98 | ON337681/ SAMN27735013 | Igarapé Aiuanã, Negro River, right bank | -64.930 | -0.620 | unpubl. data |
| C_melano_CTGAM5730 | *C. melanocephalus* | | ✓ | CTGAM5730 | | SAMN27735008 | Unknown | | | unpubl. data |
| C_melano_CTGAM5732 | *C. melanocephalus* | | ✓ | CTGAM5732 | | SAMN27735009 | Unknown | | | unpubl. data |
| C_melano_INPA5238_Amana | *C. melanocephalus* | ✓ | | INPA5238 | MvR 021 | EU560419 | Amanã Lake, Solimões | -64.500 | -2.500 | Boubli et al. 2008 |

| | | | | | | | | | | |
|---|---|---|---|---|---|---|---|---|---|---|
| | | | | | | | River, Brazil | | | |
| C_melano_INPA5239_Amana | *C. melanocephalus* | ✓ | | INPA5239 | MvR 022 | EU560420.1 | Amanã Lake, Solimões River, Brazil | -64.500 | -2.500 | Boubli et al. 2008 |
| C_melano_CTGAM5705_Serr | *C. melanocephalus* | ✓ | ✓ | CTGAM5705 | JPB110 | EU560422/ SAMN27735007 | Amanã Lake, Solimões River, Brazil | -65.170 | -0.470 | Boubli et al. 2008 |
| C_melano_FJ531640.1_Inirida | *C. melanocephalus* | ✓ | | FMNH88250 | | FJ531640.1 | Inirida River, Colômbia | -70.400 | 2.300 | Figueiredo-Ready et al. 2013 |
| C_melano_FJ531641.1_Inirida | *C. melanocephalus* | ✓ | | FMNH88251 | | FJ531641.1 | Inirida River, Colômbia | -70.400 | 2.300 | Figueiredo-Ready et al. 2013 |
| C_melano_FJ531642.1_Clbia | *C. melanocephalus* | ✓ | | FMNH89470 | | FJ531642.1 | Barracon, Alto Cano Itilla, Colômbia | -72.690 | 1.610 | Figueiredo-Ready et al. 2013 |
| C_melano_FJ531643.1_Vaupes | *C. melanocephalus* | ✓ | | FMNH89471 | | FJ531643.1 | Cano Miraflores, Vaupés River, Colômbia | -72.000 | 1.500 | Figueiredo-Ready et al. 2013 |
| C_melano_FJ531644.1_Vaupes | *C. melanocephalus* | ✓ | | FMNH89468 | | FJ531644.1 | Lago el Dorado, Vaupés River, Colômbia | -70.450 | 1.000 | Figueiredo-Ready et al. 2013 |
| C_melano_FJ531645.1_Vaupes | *C. melanocephalus* | ✓ | | FMNH89469 | | FJ531645.1 | Lago el Dorado, Vaupés River, Colômbia | -70.450 | 1.000 | Figueiredo-Ready et al. 2013 |
| C_melano_FJ531646_Mncapuru | *C. melanocephalus* | ✓ | | MVZ-016 | | FJ531646 | Manacapuru River, Brazil | -61.571 | -3.032 | Figueiredo-Ready et al. 2013 |
| C_melano_FJ531647_Mncapuru | *C. melanocephalus* | ✓ | | MVZ-017 | | FJ531647 | Manacapuru River, Brazil | -61.571 | -3.032 | Figueiredo-Ready et al. 2013 |
| Chiro_sagulatus_CTGAM515 | *Chiropotes sagulatus* | | ✓ | CTGAM515 | TRO515 | SAMN27735028 | Saracá Taquera National Forest, Trombetas River, right bank, Brazil | -56.797 | -1.488 | unpubl. data |
| Chiro_sagulatus_CTGAM674 | *Chiropotes sagulatus* | | ✓ | CTGAM674 | TRO674 | SAMN27735029 | Trombetas Biological Reserve, Trombetas River, left bank | -56.717 | -1.423 | unpubl. data |
| Chiro_sagulatus_FJ531667.1 | *Chiropotes sagulatus* | ✓ | ✓ | | | FJ531667.1 | Rio Trombetas, Pará, Brazil | | | Figueiredo-Ready et al. 2013 |
| Pit_irrorata_CTGAM426 | *Pithecia irrorata* | | ✓ | | | SAMN27735030 | unknown | | | unpubl. data |
| Chera_purinus_CTGAM209 | *Cheracebus purinus* | | ✓ | | | SAMN27735027 | Abufari Biological Reserve, left bank of Purus River, Amazonas state, Brazil | -62.960 | -4.984 | unpubl. data |
| Plect_cupreus_AAM15 | *Plecturocebus cupreus* | | ✓ | | | SAMN27735031 | Catuá-Ipixuna Extractive Reserva, Amazonas state, Brazil | -63.897 | -3.855 | unpubl. data |

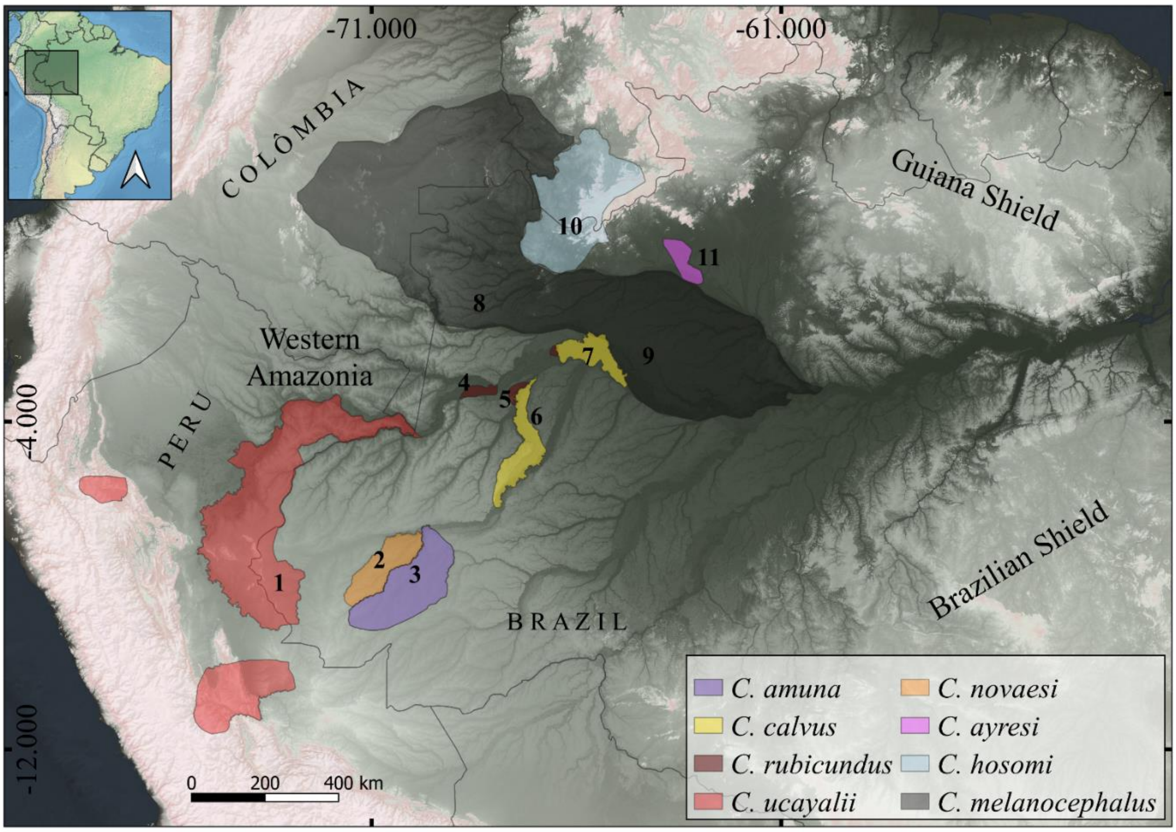


**FIGURE S1. The geographic distribution of *Cacajao* species in the central and western Amazonia. Numbers indicate the populations (see Table 2).**

**TABLE S2. Population and sample set used in this study. See also Figure S1 for population distribution.**

| Map | Species (locality) | Code | ddRAD sequences |
|---|---|---|---|
| 1 | *C. ucayalii* (Serra do Divisor National Park) | uca | 2 |
| 2 | *C. novaesi* (Gregório-Tarauacá interfluve) | nov | 5 |
| 3 | *C. amuna* (Tarauacá-Pauini interfluve) | amu | 7 |
| 4 | *C. rubicundus* (Jacurapá channel, north bank of Solimões River) | rub | 3 |
| 5 | *C. rubicundus* (Jutaí River, left bank) | rubJT | 3 |
| 6 | *C. calvus* (Jutaí River, right bank) | calJT | 5 |
| 7 | *C. calvus* (Mamirauá SDR) | cal | 3 |
| 8 | *C. melanocephalus* (upper Japurá River | melW | 4 |
| 9 | *C. melanocephalus* (Amanã SDR and Negro River) | mel | 6 |
| 10 | *C. hosomi* | hos | 2 |
| 11 | *C. ayresi* | ayr | 5 |

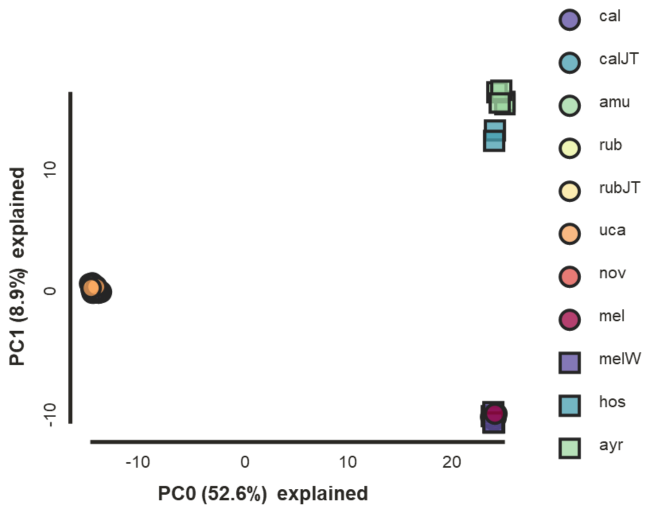


**FIGURE S2. The first two principal components of the PCA showing the allele frequency variation in *Cacajao*.** While it is possible to distinguish *C. melanocephalus*, *C. ayresi*, and *C. hosomi* clusters, the bald-headed uakaris individuals are overlapped in this graphical representation.

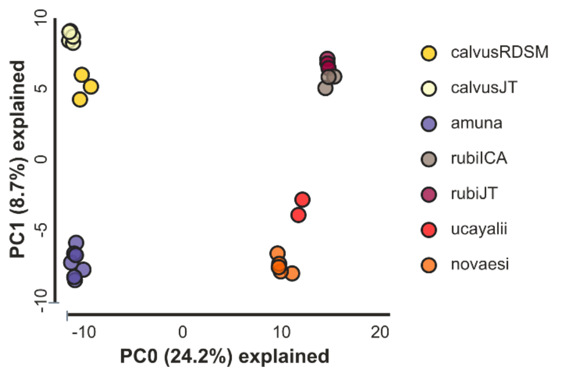


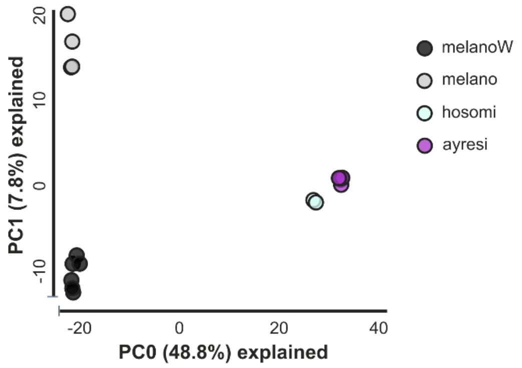


**FIGURE S3. The first two principal components of the PCA and the allele frequency variation in bald-headed uakaris (left) and in black uakaris (right).**

**Table S3. The composite likelihood and relative contribution weights for the four models testing the demographic history between *C. rubicundus*, *C. calvus*, and *C. amuna*.**

| Model | Demographic conditions | N params | Max ln (L) | AIC | AICw |
|---|---|---|---|---|---|
| Model 1 | Isolation with gene flow among all taxa | 12 | -1111.543 | 51130.845 | 0.00 |
| Model 2 | Isolation with gene flow between *C. calvus* and *C. amuna*, but no gene flow between *C. calvus* and *C. rubicundus* | 11 | -1015.465 | 4687.389 | 0.00 |
| Model 3 | Same as Model 2, but with population expansion | 18 | -997.241 | 4610,465 | 1.00 |
| Model 4 | Same as Model 2, but with a population decline | 18 | -1188.725 | 5469,281 | 0.00 |
